# Visualizing ultrastructural details of placental tissue with super-resolution structured illumination microscopy

**DOI:** 10.1101/2020.02.22.960559

**Authors:** Luis E. Villegas-Hernández, Mona Nystad, Florian Ströhl, Purusotam Basnet, Ganesh Acharya, Balpreet S. Ahluwalia

## Abstract

Super-resolution fluorescence microscopy is a widely employed technique in cell biology research, yet remains relatively unexplored in the field of histo-pathology. Here, we describe the sample preparation steps and acquisition parameters necessary to obtain fluorescent multicolor super-resolution structured illumination microscopy (SIM) images of both formalin-fixed paraffin-embedded and cryo-preserved placental tissue sections. We compare super-resolved images of chorionic villi against diffraction-limited deconvolution microscopy and demonstrate the significant contrast and resolution enhancement attainable with SIM. We show that SIM resolves ultrastructural details such as the syncytiotrophoblast’s microvilli brush border, which up until now has been only resolvable by electron microscopy.

## Introduction

Traditionally, fluorescent optical microscopy served as a tool to observe cellular dynamics with high specificity at low resolution, whereas electron microscopy helped to visualize ultrastructural details but at the cost of specificity. About two decades ago, the development of fluorescence-based super-resolution fluorescence microscopy (SRM), also known as optical nanoscopy, bridged the gap between optical and electron microscopy, enabling high specificity studies at the sub-cellular level [1]. Among the existing super-resolution methods, structured illumination microscopy (SIM) stands out as the fastest and most suitable technique for histological examination, enabling high-resolution and high-throughput imaging for decision support in clinical settings [2, 3].

In placental biology, SRM studies have focused primarily on trophoblast cell cultures [4, 5], leaving placental tissues comparatively unexplored. Here we describe the sample preparation steps and imaging parameters necessary to obtain super-resolved multicolor micrographs of both formalin-fixed paraffin-embedded (FFPE) and cryo-preserved placental sections with SIM. We report our observations of imaging chorionic villi using optical nanoscopy and compare them with the well-established diffraction-limited deconvolution microscopy (DV). To the best of our knowledge, this is the first report of SIM on placental tissue.

## Materials and Methods

Full-term placentae were collected immediately after delivery at the University Hospital of North Norway. Written informed consent was obtained from the participants according to the protocol approved by the Regional Committee for Medical and Health Research Ethics of North Norway (REK Nord reference no. 2010/2058-4).

The samples were prepared in two ways according to the preservation technique, as outlined in the orange and cyan boxes of Figure 1.

**Figure 1:**
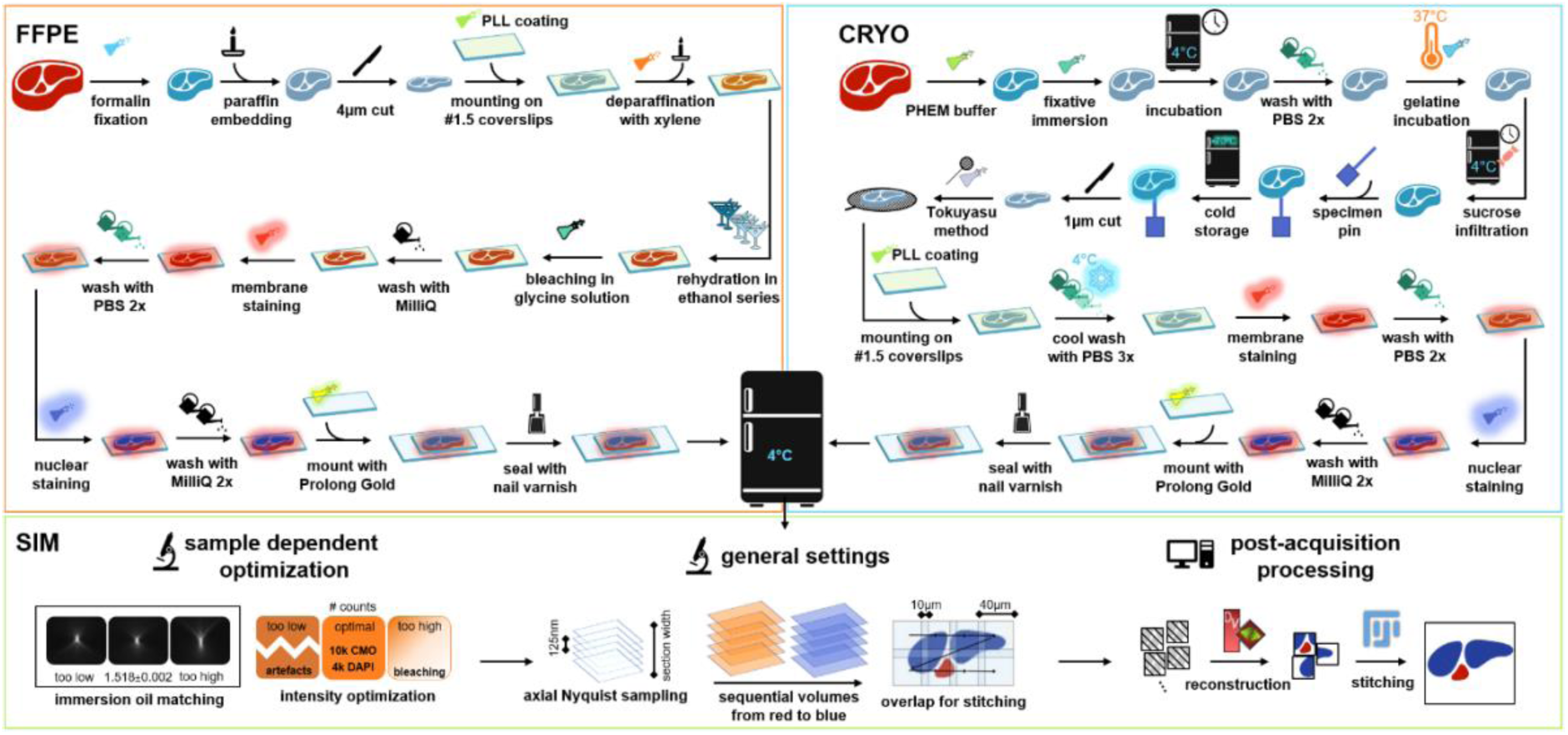
Sample preparation protocols and acquisition steps for successful SIM imaging of placental tissue sections.

### Preparation of FFPE-sections

Placental tissue samples were fixed in formalin and embedded in paraffin, according to standard histological procedures [6]. The paraffin blocks were cut into 4 μm thick sections (HM 355S Automatic Microtome, Thermo Fisher Scientific, Waltham, Massachusetts, USA), placed on poly-L-lysine coated #1.5 coverslips, and deparaffinized in xylene (3 × 5 min), followed by rehydration in descendent series of ethanol: 100% (2 × 10 min), 96% (2 × 10 min) and 70% (10 min). The rehydrated samples were immersed in bleaching solution (30 min) and then washed with MilliQ water (5 min), before incubation with CellMask Orange (CMO) membrane staining (10 min). Thereafter, the samples were washed with PBS (2 × 5 min) and incubated with DAPI nuclear staining (5 min), followed by a wash with MilliQ water (2 × 5 min). The stained samples were mounted in the center of standard microscope glass slides with Prolong Gold and sealed with nail varnish after the mounting medium hardened. All incubations were performed at room temperature. For autofluorescence characterization, the samples were prepared according to the above-described steps, excluding fluorescent dyes. Supplementary information S1 provides a detailed list of the materials used in the study.

### Preparation of cryosections

Chorionic tissue was collected, dissected and rinsed in sodium chloride as described elsewhere [7]. Collected tissue samples were transferred to 5 mL of 1×PHEM-buffer. Placental samples of 1 mm^3^ were immersed in 5 ml 8% formaldehyde in PHEM buffer and incubated at 4°C overnight. Tissue samples were immersed in 0.12% glycine at 37°C (1 h), infiltrated with 2.3 M sucrose at 4 °C overnight and transferred to specimen pins before storage in liquid Nitrogen. The samples were cut into 1 μm thick cryosections (EMUC6 ultramicrotome, Leica Microsystems, Vienna, Austria), collected with a wire loop filled with a 1:1 mixture of 2% methylcellulose and 2.3 M sucrose according to the Tokuyasu method [8], and placed on poly-L-lysine coated #1.5 coverslips. Thereafter, the samples were washed with PBS (3 × 7 min) at 4°C, followed by the staining and mounting steps previously described for the FFPE sections, at room temperature.

After preparation, both FFPE and cryosections were protected from light and stored at 4°C before imaging.

### Optical nanoscopy

The samples were imaged in SIM mode using a commercial microscope DeltaVision OMX V4 Blaze imaging system (GE Healthcare, Chicago, USA) equipped with a 60X/NA1.42 oil-immersion objective lens (Olympus, Tokyo, Japan). The acquisition parameters were optimized following the steps depicted in the green box of Figure 1. These include (1) refractive index matching of the immersion oil, (2) choice of appropriate excitation intensity and exposure time, (3) selection of sampling steps along the optical axis of the microscope, (4) sequential image acquisition from long to short wavelengths, and (5) overlap between adjacent fields of view to enable mosaic stitching of the data. After the acquisition, the super-resolved SIM images were reconstructed using the software package SoftWoRx provided by the microscope’s manufacturer, and post-processed with the open-source software Fiji [9]. Diffraction-limited deconvolution microscopy (DV) images were acquired for comparative analysis. Supplementary information S2 provides a detailed description of the acquisition parameters used in the study, as well as the working principles of SIM and DV.

## Results and Discussion

The advantages of SIM includes rejection of out-of-focus light and a two-fold resolution enhancement in all three axes (x, y, z) which gives an eight-fold contrast improvement as compared to diffraction-limited microscopy techniques such as DV. In the case of FFPE placental sections (Figure 2A,E), SIM provides a sharp visualization of biological structures such as the syncytiotrophoblasts (SYN), individual cytotrophoblasts (CT), fetal capillaries (FC), fetal red blood cells (fRBC) and maternal red blood cells (mRBC), as well as subcellular details such as the plasma membrane border (PM) between adjacent cells. In comparison, results from diffraction-limited DV microscopy offer significantly fewer details and contrast (Figure 2C,F). Notably, the resolution attainable with SIM on FFPE sections was largely limited by the level of ultrastructural preservation of the sample during the preparation steps and not by the SIM method per se. The main challenges with FFPE sections were the autofluorescence signal and the refractive index mismatch between the specimen and the imaging medium, which often led to reconstruction artifacts on the SIM images. We successfully minimized the artifacts by introducing a bleaching step in the preparation protocol and a series of iterative oil changes to match the refractive index, as illustrated in supplementary information S2 and S3.

**Figure 2:**
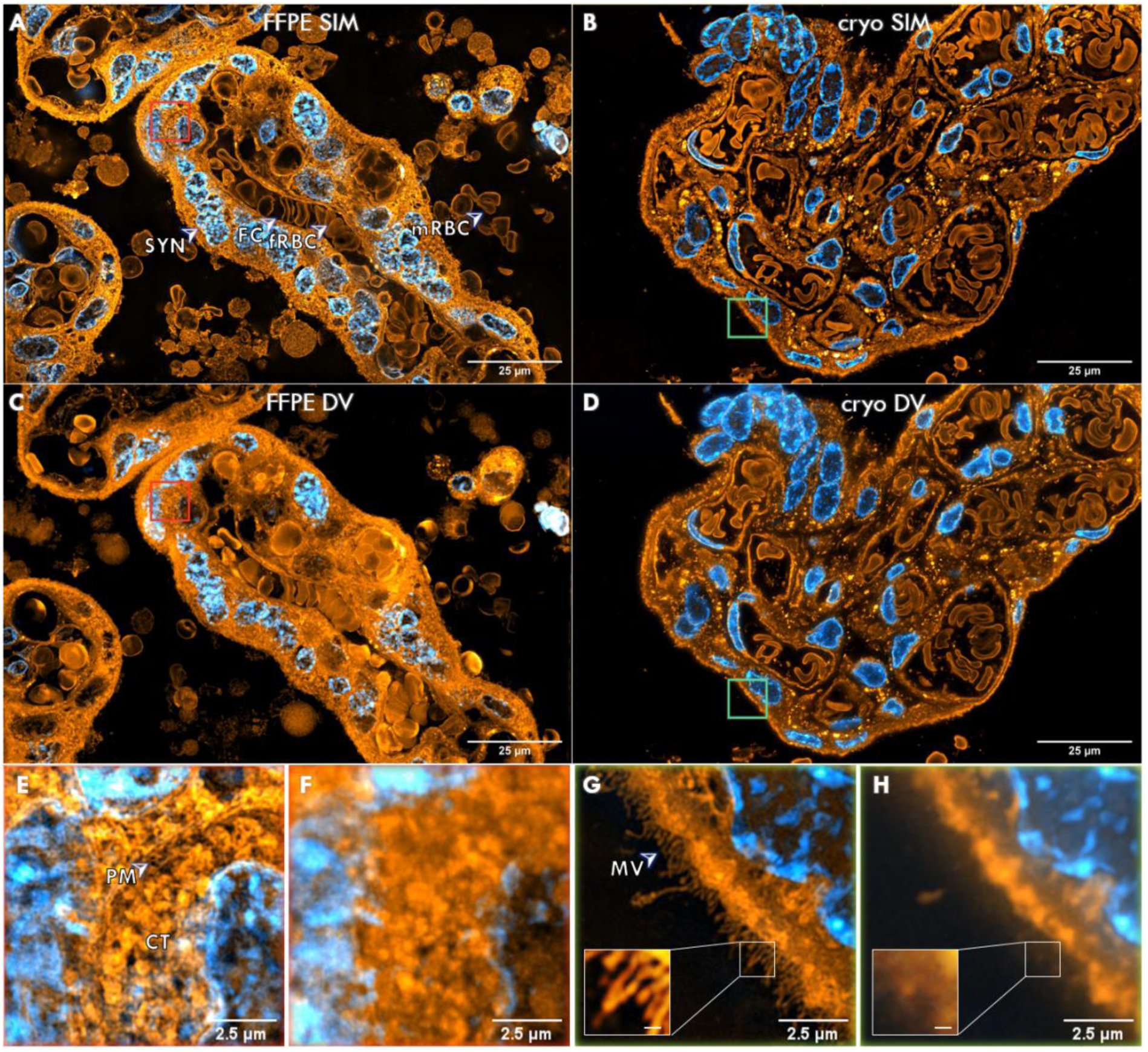
(A,C) SIM and DV images of a 4 μm thick FFPE placental section of chorionic villi. Membranes in orange (stained with CMO) and nuclei in cyan (stained with DAPI). Relevant structures such as the syncytiotrophoblasts (SYN), fetal capillaries (FC), fetal red blood cells (fRBC) and maternal red blood cells (mRBC) become observable with the enhanced contrast offered by SIM. (E,F) The enhanced resolution allows the visualization of subcellular details such as the plasma membrane (PM) of an individual cytotrophoblast (CT). (B,D) SIM and DV images of a 1 μm thick cryosection of chorionic villi. Membranes in orange (stained with CMO) and nuclei in cyan (stained with DAPI). (G,H) Magnified view of cryosections allows observation of syncytiotrophoblast microvilli (MV) brush border in SIM. Inlays have digitally enhanced contrast; scale bars are 100 nm.

The SIM images obtained from placental cryosections were significantly sharper and richer in detail than their FFPE counterparts (Figure 2B,G). The relatively low autofluorescent signal and the thin section thickness of these samples further aided the contrast enhancement of SIM by reducing the out-of-focus information. Moreover, the ultrastructural preservation attainable with the Tokuyasu method allowed the visualization of ultrastructural details such as the syncytiotrophoblast’s microvilli brush border (MV), which up until now was only resolvable by EM techniques [10].

A further advantage of imaging cryosections is the ease of tissue preparation, which allowed for sectioning, staining, and imaging within the same day. This positions SIM very favorably over traditional techniques such as EM, where the sample preparation alone can take several days to weeks. The main drawback of imaging cryosections is the dependence of high precision cutting instruments together with highly skilled operators to obtain sections with minimal morphological damage.

Taken together, the availability of optical super-resolution microscopes in conjunction with optimized tissue preparation and imaging parameters is likely to simplify and speed up the imaging of placental tissue samples aiding to better identification of abnormal structures and lesions which can be expected to improve the quality of histopathological diagnosis of placental disorders and improve image-based research in placental biology.

## Acknowledgments

We thank Randi Olsen for cryosectioning and Mona Pedersen for making FFPE tissue sections. BSA acknowledges the funding from the Research Council of Norway, (project # NANO 2021 – 288565), (project # BIOTEK 2021 – 285571). FS acknowledges funding from a Horizon2020 Marie Skłodowska-Curie Action (836355).

## Supplementary information

### S1. Materials used in the study

Table S1 and Table S2 provide a detailed list of the materials, reagents, dyes and mounting medium used in the study.

**Table S1.** Materials, reagents, dyes and mounting medium used in the study

| Material/<br>reagent | Manufacturer | Catalog<br>number | Stock<br>concentration | Working<br>concentration | Purpose |
| --- | --- | --- | --- | --- | --- |
| #1.5 coverslip | VWR | 48393-151 | - | - | Coverslip, sample substrate |
| Poly-L-lysine | Sigma-Aldrich | P8920 | 0.1 % (w/v) in H<sub>2</sub>O | 1:1 | Surface coating |
| Xylene | VWR | 28975 | ≥98.5% | 1:1 | Deparaffinization steps |
| Ethanol (EtOH) | Sigma-Aldrich | 32205-M | ≥99.8% | 100%<br>96%<br>70% | Rehydration of paraffin-embedded samples |
| Glycine | Sigma-Aldrich | G7126 | ≥99% | 150 mM | Bleaching solution to reduce autofluorescence |
| CellMask Orange | ThermoFisher | C10045 | 5 mg/mL | 1:2000 | Membrane staining |
| DAPI | ThermoFisher | 62247 | 1 mg/mL | 1:1000 | Nuclear staining |
| Prolong Gold | ThermoFisher | P36930 | - | 1:1 | Mounting medium |
| Phosphate buffered saline (PBS) | Sigma-Aldrich | D8662 | - | 1:1 | Washing steps |
| Sodium chloride | Fresenius Kabi | 826968 | 9 mg/mL | 1:1 | Rinsing steps for cryosections |
| Methyl cellulose | Sigma-Aldrich | M6385 | - | 2% | Preservation steps of cryosections |
| Sucrose | Sigma-Aldrich | 16104 | - | 2.3 M | Preservation steps of cryosections |
| Fixative | 15 mL 0.2 M PBS<br>7.5 mL 8 % formaldehyde diluted in 4× PHEM<br>7.5 mL ddH<sub>2</sub>O |  |  |  | Fixation of cryosections |

**Table S2.** Chemicals used to make the 4× PIPES-HEPES-EGTA-Magnesium sulfate (PHEM) buffer

| Chemicals | Manufacturer | Reference | Purpose |
| --- | --- | --- | --- |
| EGTA<sup>1</sup> | Sigma-Aldrich | E4378-100G | Used in PHEM |
| HEPES<sup>2</sup> | VWR Chemicals | 441476L | Used in PHEM |
| PIPES<sup>3</sup> | Sigma | P6757-500G | Used in PHEM |
| Magnesium sulfate | Sigma-Aldrich | M7506-500G | Used in PHEM |
| 5 M sodium hydroxide | Sigma-Aldrich | 30620-1KG-R | Used in PHEM |

<sup>1</sup> Ethylene glucol tetraacetic acid (ethylene glycol-bis (β-aminoethyl ether)-N,N,N',N'-tetraacetic acid) (EGTA)
<sup>2</sup> 4-(2-hydroxyethyl)-1-piperazineethanesulfonic acid (HEPES)
<sup>3</sup> 1,4 piperazine bis (2-ethanesulfonic acid) (PIPES)

### S2. Imaging parameters

#### The OMX microscope

The samples were imaged in both structured illumination microscopy (SIM) mode and deconvolution microscopy (DV) mode using a commercial microscope DeltaVision OMX V4 Blaze imaging system (GE Healthcare, Chicago, USA). The system was equipped with a 60X/NA1.42 oil-immersion objective lens (Olympus, Tokyo, Japan), three sCMOS cameras, four excitation lasers (405 nm, 488 nm, 568 nm, 642 nm), four emission filters (419-450nm, 504-552 nm, 590-627 nm, 663-703 nm). Unless otherwise stated, only the 568 nm channel for CellMask Orange (CMO) and the 405 nm channel for nuclear stains (DAPI) were used.

Super-resolved SIM images were reconstructed using the software package SoftWoRx provided by the microscope’s manufacturer, and post-processed with the open-source software Fiji to allow stitching of larger regions. Diffraction-limited deconvolution microscopy (DV) images were acquired for comparative analysis. The working principle of these microscopy techniques is hereby described.

#### Working principle of deconvolution microscopy (DV)

DV is a diffraction-limited technique used in fluorescence microscopy to enhance image contrast by removing out-of-focus information from adjacent planes above and below a given focal plane. The method involves the acquisition of multiple images along the optical axis of the microscope (e.g., z-stack), which are enhanced computationally in a post-processing step. During this step, following a mathematical model of diffraction and imaging noise, out-of-plane diffracted light is reassigned to its original location [11]. DV offers high-resolution fluorescent images near the optical resolution limit of the microscope, comparable to confocal microscopy.

#### Working principle of structured illumination microscopy (SIM)

SIM is a fluorescent-based optical super-resolution technique that employs multiple laser beams interference to create a striped illumination pattern with high spatial frequency for the excitation of the fluorescent markers on the sample. Upon illumination, the very fine, conventionally unresolvable details of the sample become down-modulated, generating coarse-spaced Moiré fringes that are now resolvable by the microscope. To computationally extract the down-modulated details and obtain super-resolved information from the sample, a set of individual images are taken over the same field of view while varying the phase and the orientation of the illumination pattern. Typically, 15 images are used for 3D-SIM to gain an isotropic lateral resolution enhancement [12], corresponding to three orientations and five phases per orientation angle. Finally, a computational algorithm processes all the raw images and reconstructs a single super-resolved SIM image.

Mathematically, the resolution limit *R* of the microscope in the spatial domain is denoted as the cut-off frequency *f*_*c*_ in the frequency domain, and their relationship is given by equation (1). Hence, the objective lens can be described as a low-pass filter in the frequency domain with a characteristic cut-off frequency *f*_*c*_.

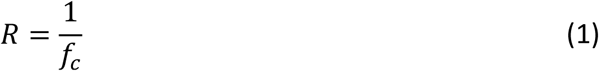

For SIM, the resolution limit *R*_*SIM*_ is given by equation (2), where *f*_*s*_ is the spatial frequency of the illumination pattern projected onto the sample [13].

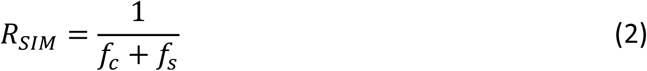

Theoretically, the resolution of SIM becomes unlimited by increasing *f*_*c*_ and *f*_*s*_. In practice, when using the same objective lens to both generate the illumination pattern and to observe the sample, both *f*_*c*_ and *f*_*s*_ are limited by the diffraction of light. Thus, the practical resolution improvement of this technique is a factor of two-fold (2X) compared to diffraction-limited microscopy, in cases where the objective lens is used both for illumination and collection of light. In practical terms, it means a resolution limit is close to 100 nm.

#### Oil matching

Both DV and SIM require optimization of the point spread function (PSF) of the system by matching the refractive index of the immersion oil to that of the mounting medium. This is an iterative process done by observing the orthogonal view of single emitters along with fluorescent images arranged in a z-stack fashion, and changing the immersion oil to a different refractive index until a symmetric shape of the PSF is observed. The PSF optimization plays a key role in the quality of DV and SIM since the oil mismatch can introduce artifacts in the reconstructed images [11, 14].

In cryosections, we matched the oil using fluorescent z-stack raw images acquired in DV mode at 250 nm sampling steps along the optical axis, as a preliminary step for SIM imaging. In FFPE sections single emitters were scarce. Instead, we imaged areas of no clinical relevance in SIM mode with different oils until artifact-free SIM images were obtained. Once optimized, the microscope’s stage was moved to the region of interest for the acquisition of SIM images. Typical refractive indices of the oils used range from 1.516 to 1.520.

#### Image acquisition

SIM images consist of a projection of multiple images (z-stack) acquired in controlled steps (z-sections, or optical sections) along the optical axis of the microscope. Following the Nyquist sampling theorem, steps of 125 nm size are required to attain SIM images of maximum 250 nm axial resolution. Projected SIM images, commonly referred to as 3D-SIM images, require 15 two-dimensional raw frames per image plane of the z-stack, in correspondence to the three angles and five phase shifts of the illumination pattern used in this technique. The thickness of the z-stack is set according to the number of optical sections needed to cover the physical thickness of the tissue section. Analogously, in the case of deconvolution microscopy (DV), only one frame is required per optical section, with a coarser sampling step of 250 nm along the optical axis.

For multi-color imaging, we sequentially acquired z-stacks of individual channels, transitioning from longer to shorter excitation wavelengths. We used two excitation laser sources for SIM imaging, namely 568 nm for membranes (CMO) and 405 nm for nuclei (DAPI). Imaging parameters such as illumination intensity and exposure time were optimized during the imaging process to avoid photobleaching of the dyes. The maximum intensity count was approximately 10000 for CMO and approximately 4000 for DAPI. The field of view (FOV) of reconstructed 3D-SIM images was approximately 40 × 40 μm, corresponding to an image size of 1024 × 1024 pixels. For larger FOVs, the samples were scanned at lateral steps of 30 μm, i.e. with 10 μm overlap, and the collected images were computationally stitched as tile mosaic images using the grid/collection stitching plugin provided by the software package FIJI [15].

### S3. Autofluorescence quantification of placental FFPE sections

One of the main challenges faced during the study was the autofluorescent background. Autofluorescence is a phenomenon that refers to the ability of certain molecular structures to fluoresce naturally or due to chemical changes induced by fixation. Autofluorescence often exhibits a broad emission spectrum, making it difficult to separate its signal from that of fluorescent dyes linked to target structures in imaging experiments. Signal contribution from autofluorescence leads to reconstruction artifacts in SIM such as haloing and high-frequency noise, thus hampering the resolution capabilities of this technique. The effect is even more prominent in conventional fluorescence microscopy (e.g. DV), where autofluorescent signal increases the background noise, leading to image blurring.

We found autofluorescence mostly in FFPE prepared sections and thus limited its characterization to this preparation method. For characterization, FFPE sections were label-free prepared and imaged in DV mode on the OMX microscope. All channels were set to 50% excitation intensity and 50 ms acquisition time. The autofluorescence response of FFPE tissue sections is illustrated in Figure S1 with mean and maximum gray values listed in Table S3. The FFPE placental tissue sections show a broad autofluorescent response with a peak on channel 3, corresponding to an excitation wavelength of 488 nm. A significant part of the autofluorescent contribution in these samples came from the hemoglobin of red blood cells (RBCs), which is known to be autofluorescent at this wavelength range [16].

Diverse methods exist to reduce autofluorescence of histological samples, ranging from pre-bleaching via irradiation [17], to a series of chemical treatments [18-20]. The chosen method for autofluorescence reduction in this study consisted of a 30 min incubation step in 150 mM glycine solution after rehydration of the tissue sections. Also, the extrinsic fluorescent dyes selected for structure labeling exhibited a signal strength at least two-fold higher than that of the autofluorescent signal in their given excitation channel.

**Figure S1.**
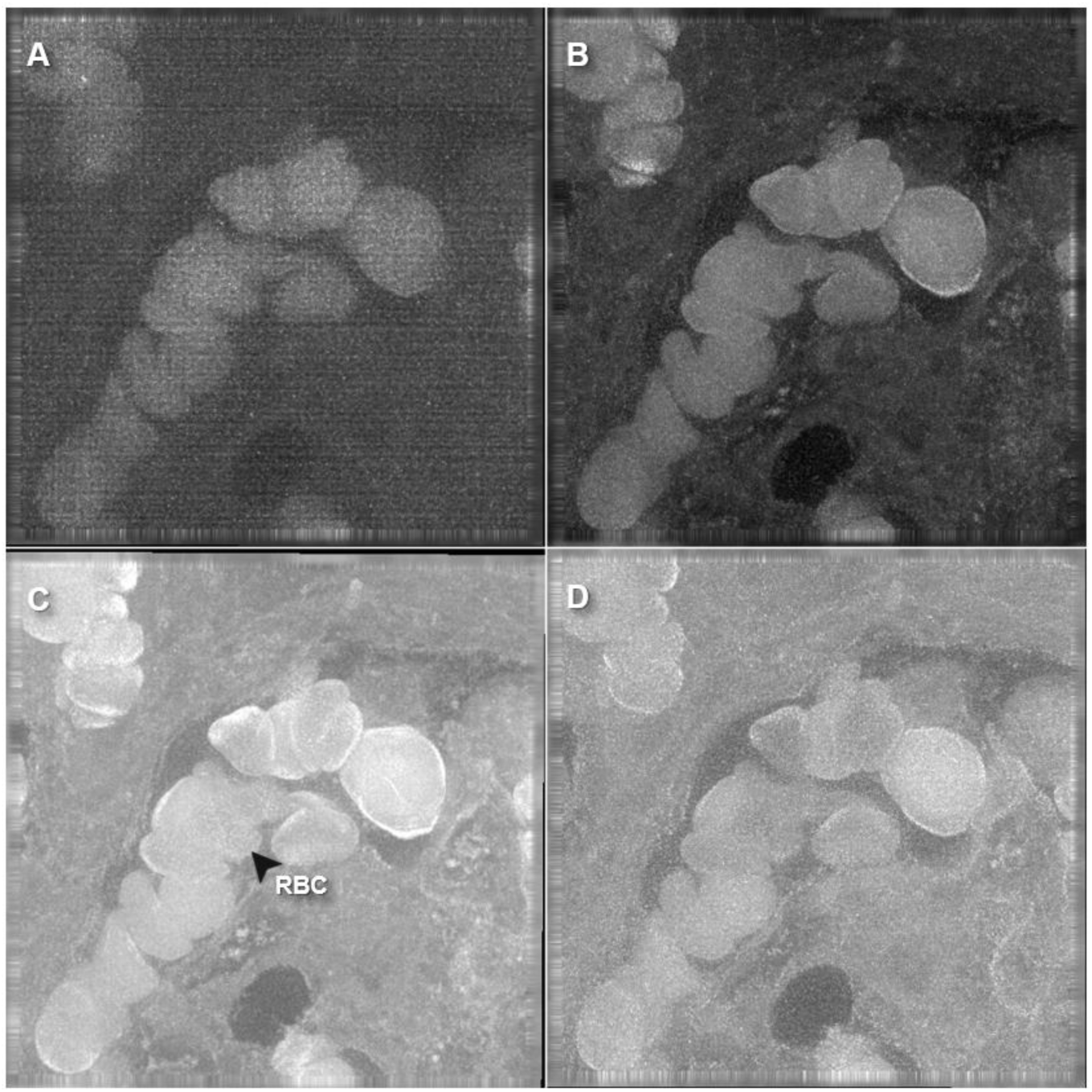
DV image of unlabeled FFPE human placental tissue section. Grayscale images show autofluorescent signal upon illumination with (a) red light (λ = 642 nm), (b) green light (λ = 568 nm), (c) blue light (λ = 488 nm), and (d) violet light (λ = 405 nm). Bright areas indicate elevated autofluorescent signal, in this case corresponding to autofluorescence of RBCs. Excitation intensity 50%, exposure time 50ms in all channels. FOV 80 × 80 μm.

**Table S3.** Imaging parameters and autofluorescence values of FFPE human placental tissue section.

| Channel | Excitation wavelength (nm) | Intensity (%) | Exposure time (ms) | Gray value count |  |  |  |
| --- | --- | --- | --- | --- | --- | --- | --- |
|  |  |  |  | Min | Max | Mean | StdDev |
| CH1 | 642 | 50 | 50 | 0 | 647 | 209 | 58 |
| CH2 | 568 | 50 | 50 | 437 | 2989 | 1151 | 299 |
| CH3 | 488 | 50 | 50 | 0 | 8596 | 4875 | 901 |
| CH4 | 405 | 50 | 50 | 0 | 2190 | 1294 | 165 |

